# Carry-over effects of larval microclimate on the transmission potential of a mosquito-borne pathogen

**DOI:** 10.1101/211037

**Authors:** Michelle V. Evans, Justine C. Shiau, Nicole Solano, Melinda A. Brindley, John M. Drake, Courtney C. Murdock

## Abstract

Climate shapes the transmission of mosquito-borne pathogens through impacts on both the vector and the pathogen. In addition to direct effects of the environment, carry-over effects from previous life history stages can influence mosquito traits relevant to disease transmission. While this has been explored in a laboratory setting, the net effect of temperature-mediated carry-over effects due to relevant environmental variation in the larval stage is ambiguous. Here, we use data collected from a semi-field experiment investigating dengue dynamics in *Aedes albopictus* across a natural environmental gradient to parameterize a dengue transmission model. We reared *Ae. albopictus* across three different land classes characterized by their proportion of impervious surface. Emerged females were offered a dengue infectious bloodmeal, kept at a constant 27 °C, and assayed for infection, dissemination, and infectiousness 21 days post infection. Incorporating carry-over effects of larval environment on measures of vector competence resulted in lower predicted dengue transmission potential across land class and season, however a strong positive relationship with larval environmental temperature remained. Given the significant impact of carry-over effects, future mechanistic models of disease transmission should include both direct and carry-over effects of environmental temperature.

## 1 Introduction

Climate plays an important role in the transmission of mosquito-borne pathogens, determining the geographic range of disease vectors and shaping transmission dynamics. Heterogeneity in environmental conditions can directly shape individual-level variation in character traits relevant to mosquito population dynamics (1), as well as pathogen transmission (2). However, in addition to these direct effects, mosquito phenotypes can be shaped indirectly by the environmental conditions experienced in previous life history stages, a phenomenon known as carry-over effects (3). Carry-over effects have been documented in a wide-range of species with complex life cycles, such as amphibians (4), migratory birds (5), and insects (6). Similarly, the mosquito life cycle is characterized by ontogenetic niche shifts, with a larval aquatic stage and an adult terrestrial stage. Following these studies, we reason that the thermal environment a mosquito experiences during its larval stage is likely to have lasting impacts on adult traits, and, ultimately, on transmission potential.

Although it has been previously demonstrated that larval environmental temperature can alter mosquito traits important for transmission, the net effect of temperature-mediated carry-over effects on overall transmission potential is ambiguous. Current models of mosquito-borne disease typically incorporate direct effects of temperature, despite evidence that carry-over effects can have large impacts on adult phenotypes (7; 8; 9). Additionally, laboratory studies designed to estimate temperature-mediated carry-over effects are often conducted across a wider range of temperatures than mosquitoes experience in the field (10), which are not easily “scaled-up” to explain transmission across a landscape when incorporated into temperature-dependent models of mosquito-borne disease (11).

We hypothesize that relevant environmental variation during the larval stage will have lasting impacts on adult traits that are relevant for mosquito population dynamics and pathogen transmission. To assess the implications of omitting carry-over effects, we used data collected from a semi-field experiment in a the *Aedes albopictus*-dengue virus (DENV) system to parameterize a mechanistic transmission model. We then compared model predictions when carry-over effects were incorporated relative to when they were excluded.

## 2 Methods

### 2.1 Semi-Field Experimental Design

To capture natural microclimate variation mosquitoes experience in the field, we chose three replicate sites (30m^2^) each of low (0-5%), intermediate (6-40%), and high (41-100%) impervious surface, representing rural, suburban, and urban land classes, respectively, that were interspersed across Athens-Clarke County, GA following methods outlined in Murdock et al. (12) (Supp. Fig. 1). Within each site, we evenly distributed four plastic trays, each containing 100 first instar *Ae. albopictus* larvae and 1L of leaf infusion. Leaf infusion was prepared as described in Murdock et al. (12). Trays were screened with a fine mesh, placed in a wire cage to deter wildlife, covered with a clear plastic vinyl to keep rainwater from entering, and were placed in full shade. We added deionized water to trays after two weeks to prevent trays from drying up and to maintain a total water volume at 1L. We placed RFID data loggers (Monarch Instruments) in vegetation next to each tray, approximately 3 feet above the ground. Data loggers recorded instantaneous temperature and relative humidity at ten minute intervals throughout the study period. Sites were visited daily from Aug. 1 to Sept. 3, 2016 (summer replicate) and Sept. 26 to Nov. 8, 2016 (fall replicate) to quantify the number of male and female mosquitoes emerging by tray per day, mosquito body size, and the proportion of mosquitoes that can transmit DENV (vector competence).

### 2.2 Dengue virus *in vitro* culturing and mosquito infections

We propagated DENV-2 virus stock (PRS 225 488) by inoculating Vero cells with a low MOI infection. Virus-containing supernatant was harvested when the cells exhibited more than 80% cytopathic effect and stored at −80 °C. We quantified viral titers of virus stock using TCID-50 assays, calculated by the Spearman-Karber method (13). When mixed 1:1 with the red blood cell mixture, the final concentration of virus in the blood meal was 3.540 × 10^6^ *TCID_50_*/mL.

Adult mosquitoes were aggregated by site and stored in reach-in incubators at 27°*C* ± 0.5°*C*, 80%±5% relative humidity, and a 12:12 hour light:dark cycle. To ensure infected mosquitoes were of a similar age, mosquitoes were pooled into cohorts of 4-6 days old in the summer and 4-9 days old in the fall (due to slower and more asynchronous emergence rates), allowed to mate, and were fed *ad libitum* with a 10% sucrose solution. Forty-eight hours prior to infection, the sucrose was replaced with deionized water, which was then removed 12-14 hours before infection. Infectious blood meals were prepared as described in Shan et al. (14) and administered to mosquitoes through a water-jacketed membrane feeder. Blood-fed mosquitoes were then maintained as described above for the duration of the experiment.

We assessed mosquitoes for infection (bodies positive for virus), dissemination (heads positive for virus), and infectiousness (saliva positive for virus) through dissections and salivation assays 21 days post infection following Tesla et al. (15). To determine infection status, we used cytopathic effect (CPE) assays to test for the presence of virus in each collected tissue (16). Individual bodies and heads were homogenized in 500 μL of DMEM and centrifuged at 2,500 rcf for 5 minutes. 200 μL of homogenate was added to Vero cells in a solution of DMEM (1% pen-strep, 5% FBS by volume) in a 24-well plate and kept at 37 °C and 5 % *CO*_2_. Salivation media was thawed, and plated on Vero cells as above. After 5 days, Vero cells were assessed for presence of DENV-2 via CPE assays. Samples were identified as positive for virus if CPE was present in the well.

### 2.3 Intrinsic growth rates (r’) and vectorial capacity (VC)

We calculated the per capita population growth rate per tray using the relationships among the number of mosquitoes emerging per day, wing size, and fecundity (Supp. Equation 1) (17). We also calculated the vectorial capacity (*VC*; Supp. Equation 3) for each site and season using a temperature-dependent mechanistic dengue model defined in Mordecai et al. (18). To estimate vectorial capacity with and without carry-over effects, we constructed two models. The model without carry-over effects used mathematically estimated values for vector competence and fecundity based on thermal response models calculated at the adult environmental temperature (27 °C) following Murdock et al. (18), while the model incorporating carry-over effects used the empirically estimated values from our study. All other parameters were the same across the two models.

### 2.4 Statistical Analysis

All analyses were conducted with respect to the female subset of the population, as they are the subpopulation responsible for disease transmission. In the case of data logger failure (N = 3), imputed means from the site were used to replace microclimate data. In the case of trays failing due to wildlife tampering (two urban and one suburban in the fall replicate), collected mosquitoes were used for infection assays, but were excluded from survival and emergence analyses. For all mixed-models, significance was assessed through Wald Chi-square tests (*α* = 0.05) and examination of 95% confidence intervals. Pearson residuals and Q-Q plots were visually inspected for normality. All mixed models were fit using the `lme4` package in *R*.

We used generalized linear mixed models (GZLMs) to explore if microclimate (i.e. mean, minimum, maximum, and daily ranges of temperature and relative humidity), the mean proportion of adult females emerging per tray, time to female emergence, female body size, per capita growth rate, metrics of vector competence, and vectorial capacity differed across land class and season. In all models, fixed effects included land class, season, and their interaction, and site was a random effect. The effect of body size on infection dynamics was also explored at the level of the individual mosquito, fitting a binomial GZLM including wing size as a fixed effect and site as a random effect.

To explore whether observed effects of land class and season were due to variation in microclimate, we assessed the effects of microclimate on each response variable. Due to extreme correlation between variables (*ρ* > 0.9), we ultimately chose one variable to represent microclimate (mean temperature) to reduce bias due to collinearity (19). Thus, we fit GZLMs to each response variable described above with mean daily temperature and site as fixed and random effects, respectively.

## 3 Results

### 3.1 Effects of land class and season on microclimate

We found that microclimate profiles differed significantly across both season and land class (Supp. Table 1, Supp. Fig. 2). All microclimate metrics differed significantly across season, except for maximum relative humidity (*z*=0.679, *p* = 0.497). In general, temperatures were warmer in the summer and on urban sites, replicating what was found in a prior study in this system (12). Relative humidity was higher in the summer than the fall, due to a drought, and lower on urban land classes than suburban and rural classes.

### 3.2 Direct and carry-over effects of land class, season, and microclimate on population growth

The total proportion of adult females emerging per tray was significantly higher in summer than fall (Table 1), but did not differ across land class (Fig. 1A). Of the 3,600 first-instar larvae placed in each season, a total of 2595 and 1128 mosquitoes emerged in the summer and fall, respectively. There was a strong positive relationship between mean daily temperature and larval survival to emergence by tray (Table 1). The mean rate of larval development per tray was significantly different between summer and fall (Fig. 1B, Table 1), with daily development rates of 0.074 ± 0.002 day^-1^ and 0.0387 ± 0.002 day^-1^, respectively. There was a significant positive relationship between temperature and larval development rate (Table 1).

**Table 1:** Model results from mixed models investigating the effect of land class, season, their interaction, and temperature on population demographics, metrics of infection, and vectorial capacity with and without the inclusion of carry-over effects.

|  | Class |  |  | Season |  |  | Class*Season |  |  | Temperature |  |  |
| --- | --- | --- | --- | --- | --- | --- | --- | --- | --- | --- | --- | --- |
| | *df* | $\chi^2$ | p-value | *df* | $\chi^2$ | p-value | *df* | $\chi^2$ | p-value | $\beta \pm s.e.$ | t-value | p-value |
| Survival | 2 | 0.0361 | 0.982 | 1 | 61.129 | **<0.001** | 2 | 5.891 | 0.053 | 0.240 (0.0297) | 8.089 | **<0.001** |
| Development Rate | 2 | 3.847 | 0.1461 | 1 | 597.51 | **<0.001** | 2 | 3.108 | 0.211 | 0.005 (0.0002) | 20.17 | **<0.001** |
| Wing Length | 2 | 0.835 | 0.6587 | 1 | 2.7937 | 0.0946 | 2 | 14.748 | **<0.001** | 0.006 (0.003) | 1.883 | 0.061 |
| Per Capita Growth (*r'*) | 2 | 0.667 | 0.717 | 1 | 219.84 | **<0.001** | 2 | 2.622 | 0.230 | 0.013 (0.001) | 14.927 | **<0.001** |
| Infection | 2 | 18.733 | **<0.001** | 1 | 12.609 | **<0.001** | 2 | 1.985 | 0.371 | -0.075 (0.0249) | -3.011 | **<0.001** |
| Dissemination | 2 | 14.208 | **<0.001** | 1 | 15.05 | **<0.001** | 2 | 0.941 | 0.625 | -0.093(0.0282) | -3.299 | **0.004** |
| Infectiousness | 2 | 1.125 | 0.57 | 1 | 3.7762 | 0.052 | 2 | 0.302 | 0.860 | 0.006 (0.0065) | 0.955 | 0.354 |
| VC (w/o carry-over) | 2 | 0.388 | 0.824 | 1 | 79.472 | **<0.001** | 2 | 0.168 | 0.920 | 3.912 (0.449) | 8.347 | **<0.001** |
| VC (w/ carry-over) | 2 | 0.588 | 0.745 | 1 | 20.631 | **<0.001** | 2 | 0.337 | 0.845 | 0.802 (0.168) | 4.690 | **<0.001** |

**Figure 1:**
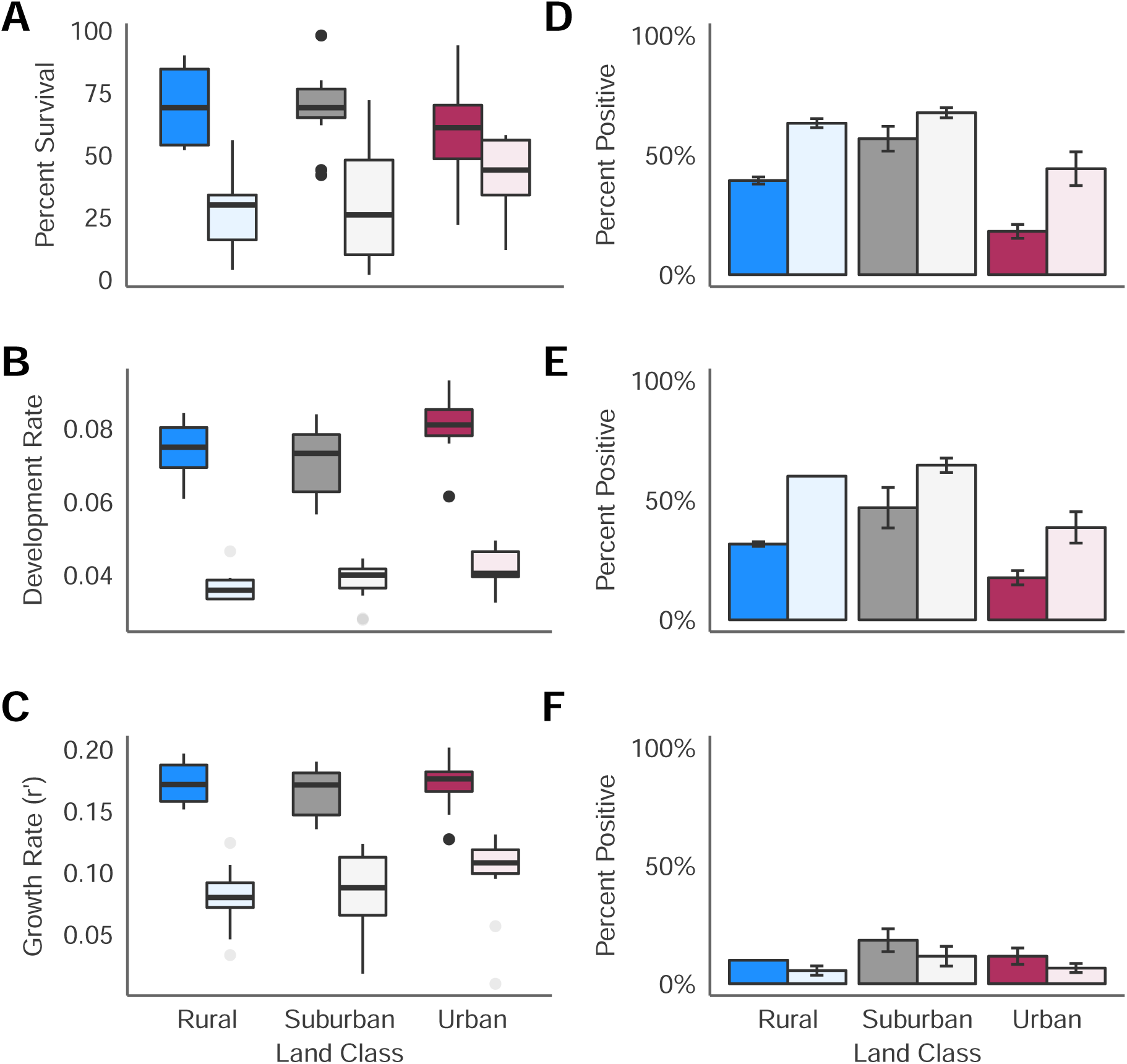
Left panel illustrates female larval a) survival rate, b) development rate and c) population growth rate across the summer (dark fill) and fall (light fill) trials and three land classes. Right panel plots mean and standard data of raw data across sites (n = 3 per treatment) of d) infection, e) dissemination, and f) infectious rates.

We did not observe a significant carry-over effect of land class or season on mosquito wing size, however there was a significant interaction between the two (Table 1), with the smallest mosquitoes on suburban and rural sites in the summer and fall, respectively. We also found no effect of temperature on female wing size. After incorporating the number of adult females emerging per day, the date of emergence, and their body size into the per capita growth rate equation (Supp. Equation 1), we found that the estimated per capita growth rate was higher in the summer season than the fall season (Fig. 1C, Table 1). There was no evidence for a difference in population growth across land class or temperature.

### 3.3 Carry-over effects of land class, season, and microclimate on vector competence

A total of 319 female mosquitoes were assessed for infection status, 20 per site in the summer and varying numbers per site in the fall due to lower emergence rates (sample sizes reported in Supp. Table 2). We found that land class and season did significantly impact the probability of a mosquito becoming infected and disseminating dengue infection (Fig. 1D, E), Table 1). The probability of becoming infectious did not differ across land class, nor season (Fig. 1F), despite the higher probability of mosquito infection and dissemination in the fall and on suburban and rural sites. This suggests that the ability of virus to penetrate the salivary glands differs in adults reared in the summer vs. the fall and across land class, with a higher proportion of dengue infected mosquitoes becoming infectious in the summer and on urban sites (Supp. Table 2, χ^2^ = 13.65, *p*< 0.001). We also found the probability of infection to decline with increasing body size (χ^2^ = 4.776, *p* = 0.0289), although there was no evidence for a relationship between body size and the probability of dissemination or infectiousness. Differences in infection status across land class and season were driven by a strong relationship with microclimate. We found that infection and dissemination rates decreased with increasing temperatures, while there was no relationship between infectiousness and temperature (Table 1).

### 3.4 Integrating direct and carry-over effects into estimates of transmission potential

When calculating *VC* with or without the inclusion of carry-over effects, *VC* was higher in the summer than the fall (Supp. Fig. 3, Table 1). In the summer season, there was a trend for *VC* to increase with increasing urbanization (Supp. Fig. 3). This trend was not significant, however, given the small sample size (n=9) and the disproportional impact of having no infectious mosquitoes at one site, resulting in a value of *VC* = 0 for one sample. Further, we found that calculated vectorial capacity increased with temperature for both models, although the increase was more pronounced when not accounting for carry-over effects (Fig. 2). When comparing *VC* calculations with and without carry-over effects, we found that including carry-over effects decreased the expected vectorial capacity overall by an average of 84.89 ± 2.86 % (Supp. Fig. 3).

**Figure 2:**
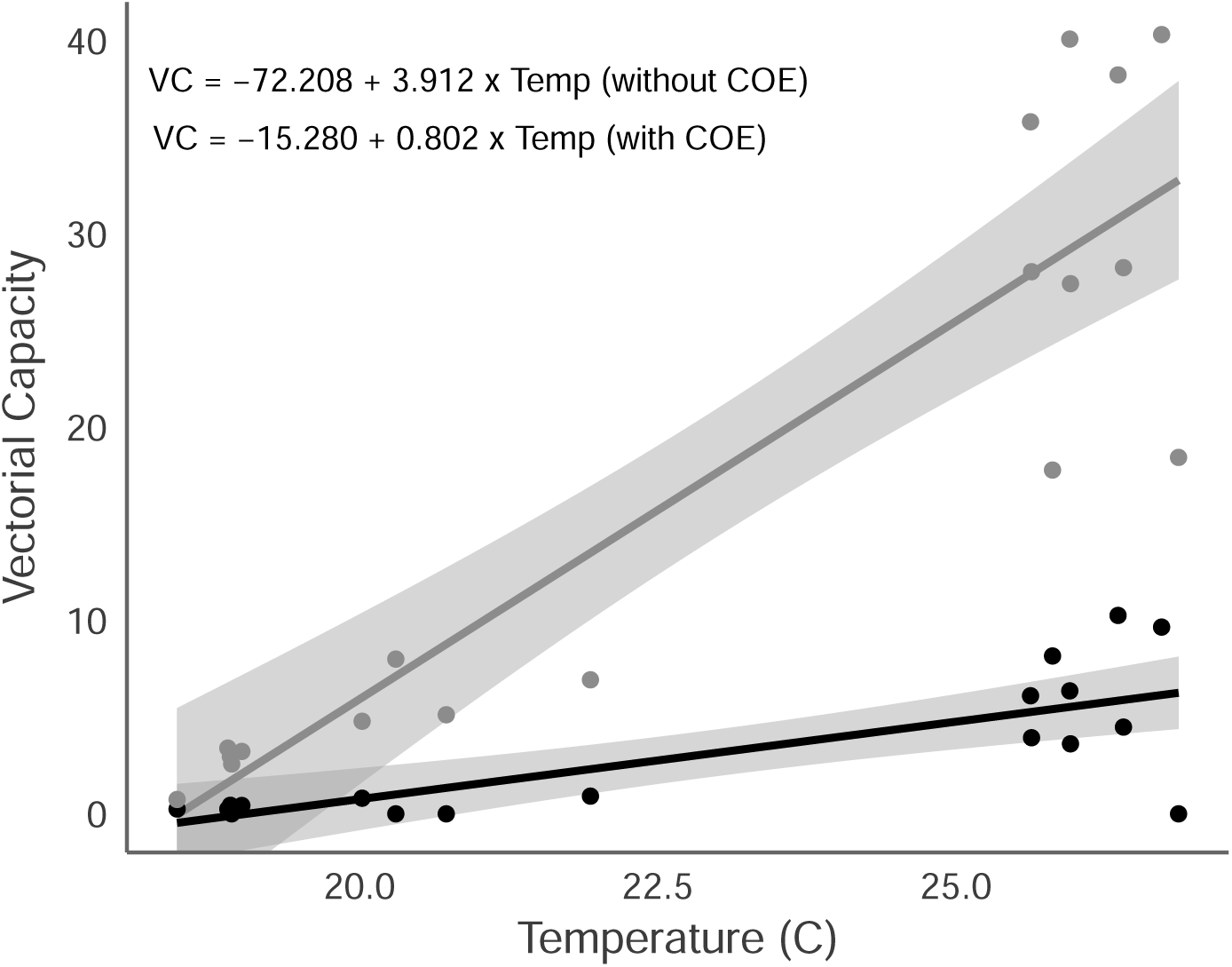
The calculated vectorial capacity (with 95 % confidence intervals) by site across individual mean temperature prior to infection assays. Light gray line represents VC without carry-over effects and dark line represent VC with carry-over effects.

## 4 Discussion

Mathematical models of mosquito-borne disease rarely include mosquito larval stages (11), and of those that do, few include the influence of carry-over effects on important mosquito life-history traits (but see (20)). This is likely because there are relatively few empirical studies parameterizing carry-over effects in mosquito-pathogen systems (21), and, most are laboratory studies conducted across a wider range of temperatures than that seen in the field. Here, we demonstrate that fine-scale differences in larval microclimate generate carry-over effects on adult vector competence and fecundity, resulting in variation in predicted mosquito population dynamics and transmission potential across an urban landscape and season.

The subtle heterogeneity in microclimate we observed resulted in significantly different predicted population growth rates through its effects on demographic traits. Daily mean temperatures (25.43 °C) across all sites in the summer were closer to the predicted thermal optimum of *Ae. albopictus* for the probability of egg to adult survival (24-25 °C) (18) than in the fall (17.69 °C), leading to higher survival rates. We also observed more rapid larval development rates in the summer relative to the fall, and on warmer urban sites in the fall only. Again, this is likely due to the strong positive relationship observed between development rates and mean larval temperature, as the metabolic rate of mosquitoes will increase with warming temperatures (1).

Surprisingly, we found no effect of land class or season on female mosquito body size, despite the difference in temperatures across season. Following allometric temperature-size relationships of ectotherms, warmer larval temperatures should lead to smaller bodied mosquitoes (22). However, our results contrast with many laboratory and field studies that have found a negative relationship between rearing temperature and mosquito body size (*Ae. albopictus* (23; 12), *Culex tarsalis* (24), *Anopheles gambiea* (25)). This may be due to a difference in nutrient quality. Nutrient availability and quality can mediate the relationship between temperature and body size (26). The majority of the above laboratory studies rear larvae on high quality food sources, such as fish food. The leaf infusion used in our experiment relied on yeast and naturally colonizing microorganisms that grow more slowly at low temperatures (27), likely constraining larval growth. For example, Lounibos et al. (28) found a positive relationship between temperature and male *Ae. albopictus* body size when larvae were provided leaf litter.

Our results agree with laboratory studies in other arboviral systems (chikungunya (29), yellow fever (29), and Rift Valley fever (30)) that found cool larval environmental temperatures to enhance arbovirus infection relative to warmer larval environments. Studies in the *Ae. albopictus*-dengue virus system have also found that low larval temperatures enhance mosquito susceptibility to viral infection, although this is dependent on larval nutrition (31) and the stage of the infection (i.e. mid-gut vs. dissemination vs. saliva) (32). While we found infection and dissemination to decrease with increasing temperatures, there was no effect of temperature on viral presence in the saliva, suggesting carry over effects due to microclimate variation may alter the overall efficiency of dengue infection. Thus, even though a smaller proportion of mosquitoes reared on urban sites and in the summer became infected and disseminated infection, these mosquitoes were more likely to become infectious, resulting in no net difference in overall vector competence across land class and season. Thus, later stages of viral infection (i.e. salivary gland penetration) may be differentially impacted by larval environmental temperature than earlier stages (i.e. midgut escape).

Current models of vector-borne disease focus primarily on direct effects of environmental variables on mosquito densities and disease transmission and rarely include the effects of the larval stage, either directly or via carry-over effects (11). We find that when carry-over effects are not incorporated, mechanistic models overestimate the effects of key environmental drivers (e.g. temperature) on transmission. The relatively small differences in temperature across our study site (less than 1.5 °C) resulted in a two-fold difference in predicted vectorial capacity when omitting carry-over effects. Thus, we would expect these phenomena to have an even larger impact in more urbanized areas, particularly megacities, with larger seasonal and spatial microclimate ranges (33).

Carry-over effects are not simply limited to microclimate, and can result due to variation in larval nutrition (34), intra- and inter-specific densities (35), and predation (20) in mosquito systems. Further, abiotic and biotic factors will likely interact to influence carry over effects (31; 36), and this interaction could be scale-dependent (37). For example, biotic processes are predicted to be more important at local geographic scales, while abiotic processes dominate at regional geographic scales in species distribution models (38). Future exploration of the scale-dependent contribution of different environmental factors and their interactive influence on both direct and carry-over effects is needed to improve models of mosquito distributions, population dynamics, and disease transmission.

In conclusion, we found fine-scale variation in microclimate to shape mosquito population dynamics and arbovirus transmission potential through direct effects on larval survival and development rates, and indirectly through carry-over effects on vector competence and fecundity. Given the devastating impact of disease in other species with complex life histories (e.g. chytridiomycosis in amphibians), the role of carry-over effects in disease transmission are an important, though understudied, mechanism that must be better understood to control disease spread. Thus, incorporating relationships between carry-over effects and organismal life-history traits into statistical and mechanistic models will lead to more accurate predictions on the distributions of species, population dynamics, and the transmission of pathogens and parasites. The interaction between the larval and adult environments, mediated by carry-over effects, could have complex consequences for adult phenotypes and fitness for mosquitoes as well as other organisms.

## 5 Acknowledgements

We thank members of the Murdock and Brindley labs for discussion and technical support conducting viral assays. We thank Diana Diaz, Abigail Lecroy, and Marco Notarangelo for assistance in the field and lab. This work was supported by the University of Georgia (Presidential Fellowship, College of Veterinary Medicine, Department of Infectious Diseases) the National Science Foundation Graduate Research Fellowship, and the National Science Foundation Research Experiences for Undergraduates (Grant No. 1156707).

## 6 Contributions

M.V.E, J.M.D, and C.C.M designed the study; M.V.E, J.C.S. and N.S. collected the data; M.V.E. and M.A.B. cultured virus and conducted CPE assays; M.V.E, J.M.D, and C.C.M analyzed the data; M.V.E., J.M.D, and C.C.M prepared the tables and figures; M.V.E., J.C.S., N.S., M.A.B., J.M.D, and C.C.M drafted the manuscript. All authors gave final approval for publication.

## 7 Data accessibility

All data and code used in analyses are available on figshare (doi:10.6084/m9.figshare.5558128).

## References

[1] Delatte H, Gimonneau G, Triboire A, Fontenille D. Influence of Temperature on Immature Development, Survival, Longevity, Fecundity, and Gonotrophic Cycles of Aedes Albopictus, Vector of Chikungunya and Dengue in the Indian Ocean. J Med Entomol. 2009 Jan;46(1):33–41.

[2] Murdock C, Paaijmans K, Bell A, King J, Hillyer J, F Read A, et al. Complex effects of temperature on mosquito immune function. Proceedings Biological sciences / The Royal Society. 2012 May;279:3357–66.

[3] Harrison XA, Blount JD, Inger R, Norris DR, Bearhop S. Carry-over Effects as Drivers of Fitness Differences in Animals. J Anim Ecol. 2011 Jan;80(1):4–18.

[4] Vonesh JR. Sequential Predator Effects across Three Life Stages of the African Tree Frog, Hyperolius Spinigularis. Oecologia. 2005 Mar;143(2):280–290.

[5] Norris DR, Taylor CM. Predicting the Consequences of Carry-over Effects for Migratory Populations. Biology Letters. 2006 Mar;2(1):148–151.

[6] De Block M, Stoks R. Fitness Effects from Egg to Reproduction: Bridging the Life History Transition. Ecology. 2005;86(1):185–197.

[7] Muturi EJ, Lampman R, Costanzo K, Alto BW. Effect of Temperature and Insecticide Stress on Life-History Traits of Culex Restuans and Aedes Albopictus (Diptera: Culicidae). J Med Entomol. 2011 Mar;48(2):243–250.

[8] Muturi EJ, Kim CH, Alto BW, Berenbaum MR, Schuler MA. Larval Environmental Stress Alters Aedes Aegypti Competence for Sindbis Virus. Trop Med Int Health. 2011 May;16(8):955–964.

[9] Price DP, Schilkey FD, Ulanov A, Hansen IA. Small Mosquitoes, Large Implications: Crowding and Starvation Affects Gene Expression and Nutrient Accumulation in Aedes Aegypti. Parasites & Vectors. 2015;8:252.

[10] Cator LJ, Thomas S, Paaijmans KP, Ravishankaran S, Justin JA, Mathai MT, et al. Characterizing Microclimate in Urban Malaria Transmission Settings: A Case Study from Chennai, India. Malaria Journal. 2013 Mar;12(1):1–1.

[11] Reiner RC, Perkins TA, Barker CM, Niu T, Chaves LF, Ellis AM, et al. A Systematic Review of Mathematical Models of Mosquito-Borne Pathogen Transmission: 1970-2010. J R Soc Interface. 2013 Apr;10(81):20120921.

[12] Murdock CC, Evans MV, McClanahan TD, Miazgowicz KL, Tesla B. Fine-Scale Variation in Microclimate across an Urban Landscape Shapes Variation in Mosquito Population Dynamics and the Potential of Aedes Albopictus to Transmit Arboviral Disease. PLOS Neglected Tropical Diseases. 2017 May;11(5):e0005640.

[13] Shao Q, Herrlinger S, Yang SL, Lai F, Moore JM, Brindley MA, et al. Zika Virus Infection Disrupts Neurovascular Development and Results in Postnatal Microcephaly with Brain Damage. Development. 2016 Nov;143(22):4127–4136.

[14] Shan C, Xie X, Muruato AE, Rossi SL, Roundy CM, Azar SR, et al. An Infectious cDNA Clone of Zika Virus to Study Viral Virulence, Mosquito Transmission, and Antiviral Inhibitors. Cell Host and Microbe. 2016 May;p. 1–23.

[15] Tesla B, Demakovsky LR, Packiam HS, Mordecai EA, Rodriguez AD, Bonds MH, et al. Estimating the effects of variation in viremia on mosquito susceptibility, infectiousness, and RO of Zika in Aedes aegypti. bioRxiv. 2017 Nov;p. 221572.

[16] Balaya S, Paul S, D’Lima L, Pavri K. Investigations on an Outbreak of Dengue in Delhi in 1967. Indian J Med Res. 1969;57(4):767–774.

[17] Livdahl TP, Sugihara G. Non-Linear Interactions of Populations and the Importance of Estimating Per Capita Rates of Change. The Journal of Animal Ecology. 1984 Jun;53(2):573–580.

[18] Mordecai EA, Cohen JM, Evans MV, Gudapati P, Johnson LR, Lippi CA, et al. Detecting the Impact of Temperature on Transmission of Zika, Dengue, and Chikungunya Using Mechanistic Models. PLOS Neglected Tropical Diseases. 2017 Apr;11(4):e0005568.

[19] Graham MH. Confronting Multicollinearity in Ecological Multiple Regression. Ecology. 2003 Nov;84(11):2809–2815.

[20] Roux O, Vantaux A, Roche B, Yameogo KB, Dabire KR, Diabate A, et al. Evidence for Carry-over Effects of Predator Exposure on Pathogen Transmission Potential. Proc R Soc B. 2015 Dec;282(1821):2015–2430.

[21] Parham PE, Waldock J, Christophides GK, Hemming D, Agusto F, Evans KJ, et al. Climate, Environmental and Socio-Economic Change: Weighing up the Balance in Vector-Borne Disease Transmission. Philosophical Transactions of the Royal Society B: Biological Sciences. 2015 Feb;370(1665):20130551–20130551.

[22] Angilleta MJ, Steury TD, Sears MW. Temperature, Growth Rate, and Body Size in Ectotherms: Fitting Peice of a Life-History Puzzle. Integrative and Comparative Biology. 2004 Jan;44:498–509.

[23] Reiskind MH, Zarrabi AA. Is Bigger Really Bigger? Differential Responses to Temperature in Measures of Body Size of the Mosquito, Aedes Albopictus. Journal of Insect Physiology. 2012 Jul;58(7):911–917.

[24] Dodson BL, Kramer LD, Rasgon JL. Effects of Larval Rearing Temperature on Immature Development and West Nile Virus Vector Competence of Culex Tarsalis. Parasites & Vectors. 2012 Sep;5(1):199.

[25] Koella JC, Lyimo EO. Variability in the Relationship between Weight and Wing Length of Anopheles Gambiae (Diptera: Culicidae). Journal of Medical Entomology. 1996 Mar;33(2):261–264.

[26] Farjana T, Tuno N, Higa Y. Effects of Temperature and Diet on Development and Interspecies Competition in Aedes Aegypti and Aedes Albopictus. Medical and Veterinary Entomology. 2011 Jul;26(2):210–217.

[27] Ratkowsky DA, Olley J, McMeekin TA, Ball A. Relationship between Temperature and Growth Rate of Bacterial Cultures. J Bacteriol. 1982 Jan;149(1):1–5.

[28] Lounibos LP, Suarez S, Menendez Z, Nishimura N, Escher RL, O’Connell SM, et al. Does Temperature Affect the Outcome of Larval Competition between Aedes Aegypti and Aedes Albopictus? J of Vec Eco. 2002 Jun;27(1):86–95.

[29] Adelman ZN, Anderson MAE, Wiley MR, Murreddu MG, Samuel GH, Morazzani EM, et al. Cooler Temperatures Destabilize RNA Interference and Increase Susceptibility of Disease Vector Mosquitoes to Viral Infection. PLOS Neglected Tropical Diseases. 2013 May;7(5):e2239.

[30] Turell M. Effect of Environmental Temperature on the Vector Competence of Aedes Taeniorhynchus for Rift Valley Fever and Venezuelan Equine Encephalitis Viruses. Am J Trop Med Hyg. 1993;49(6):672–676.

[31] Buckner EA, Alto BW, Lounibos LP. Larval Temperature-Food Effects on Adult Mosquito Infection and Vertical Transmission of Dengue-1 Virus. J Med Entomol. 2016 Jan;53(1):91–98.

[32] Alto BW, Bettinardi D. Temperature and Dengue Virus Infection in Mosquitoes: Independent Effects on the Immature and Adult Stages. The American Journal Of Tropical Medicine And Hygiene. 2013 Mar;88(3):497–505.

[33] Peng S, Piao S, Ciais P, Friedlingstein P, Ottle C, Breon FM, et al. Surface Urban Heat Island Across 419 Global Big Cities. Environ Sci Technol. 2012 Jan;46(2):696–703.

[34] Moller-Jacobs LL, Murdock CC, Thomas MB. Capacity of Mosquitoes to Transmit Malaria Depends on Larval Environment. Parasites & Vectors. 2014;7:593.

[35] Alto BW, Lounibos LP, Higgs S, Juliano SA. Larval Competition Differentially Affects Arbovirus Infection in Aedes Mosquitoes. Ecology. 2005;86(12):3279–3288.

[36] Muturi EJ, Blackshear M, Montgomery A. Temperature and Density-Dependent Effects of Larval Environment on Aedes Aegypti Competence for an Alphavirus. J Vector Ecol. 2012 Jun;37(1):154–161.

[37] Leisnham PT, LaDeau SL, Juliano SA. Spatial and Temporal Habitat Segregation of Mosquitoes in Urban Florida. PLoS ONE. 2014 Jan;9(3).

[38] Cohen JM, Civitello DJ, Brace AJ, Feichtinger EM, Ortega CN, Richardson JC, et al. Spatial Scale Modulates the Strength of Ecological Processes Driving Disease Distributions. PNAS. 2016 Jun;113(24):E3359–E3364.

